# Comparative Genomics of Two *Chaetoceros muelleri* Strains Reveals Structural and Functional Variation

**DOI:** 10.64898/2026.04.27.720831

**Authors:** Anushree Sanyal

**Affiliations:** School of Natural Sciences, Technology and Environmental Studies, Södertörn University, Sweden

## Abstract

Diatoms are major contributors to global primary production and biogeochemical cycling, yet high quality nuclear genome resources remain limited for ecologically dominant lineages such as *Chaetoceros*. Here, we present a highly contiguous nuclear genome assembly of *Chaetoceros muelleri* generated from living cells resurrected from resting spores preserved in Baltic Sea sediments and sequenced using PacBio HiFi long read technology. The resurrected BS20 assembly is highly contiguous and gapless, enabling robust resolution of genes, protein domain architecture, and transposable element (TE) landscapes that are poorly captured in fragmented assemblies.

By directly comparing this resurrected genome with a contemporary laboratory strain (*C. muelleri* NMCA1316), we disentangle biological divergence from assembly-driven artefacts. Despite a strongly conserved core genome, the two strains differ markedly in repeat content, gene family copy number, and functional enrichment patterns. The resurrected genome harbors ∼38% repetitive DNA approximately 1.8-fold higher than the laboratory strain dominated by LTR retrotransposons and a large fraction of unclassified repeats, indicating extensive historical TE activity. Expanded orthogroups in the resurrected strain are enriched for retroelement-associated domains, nucleic-acid processing functions, and stress-responsive gene families, including small heat shock proteins, whereas apparent expansions in the laboratory strain largely reflect annotation inflation arising from assembly fragmentation.

Spatial analyses further reveal widespread proximity between TEs, and genes involved in membrane transport and environmental responsiveness, suggesting a role for TE dynamics in regulatory and functional diversification. Together, our results demonstrate that assembly strategy and strain history critically shape genomic inference and highlight the value of resurrected genomes for accessing historical diversity. This study provides a foundational genomic resource for *C. muelleri* and establishes a genomic framework that explicitly accounts for strain level variation when investigating diatom genome evolution, TE mediated innovation, and long-term ecological adaptation.

## Introduction

Diatoms are a diverse and ecologically important group of microalgae that contribute nearly 20% of global primary productivity and play critical roles in marine and freshwater biogeochemical cycles (Field et al. 1998, Falkowski et al. 1998, Armbrust 2009; Tréguer et al. 2018). As major contributors to carbon fixation and silica cycling, they influence nutrient fluxes, food web dynamics, and global carbon sequestration (Smetacek 1999; Tréguer et al. 2018). The diversity of diatoms is extensive, with thousands of described species exhibiting extensive morphological, physiological, and genomic variation (Mann 1999; Kooistra et al. 2007).

Within diatoms, species of the genus *Chaetoceros* are especially widespread and ecologically important as they frequently form large blooms in coastal and offshore environments (Smayda 1997). *Chaetoceros muelleri*, a small centric diatom, has become a valuable model for studying physiological flexibility, ecological tolerance, and algal biotechnology applications due to its rapid growth and ability to thrive under diverse environmental conditions (Reynolds 2006) and for long time periods (Sanyal et al. 2022, Bolius et al. 2025). Despite its ecological relevance and application in aquaculture (Kumaran et al. 2021), genomic resources for *C. muelleri* have been limited, hindering deeper insight into its evolutionary history, metabolic potential, and environmental adaptability.

Recent advances in long-read sequencing technologies have facilitated the generation of more complete and contiguous algal genomes, improving our ability to characterize gene structure, repetitive elements, and genome organization (Armbrust et al. 2004, Bowler et al. 2008, Filloramo et al. 2021, Di Costanzo et al. 2025, Liu et al. 2024, Sorokina et al. 2022, Ogura et al. 2018). However, assemblies can differ substantially depending on sequencing depth, read type, library preparation, and computational strategy. Such differences can have downstream consequences for gene prediction accuracy, identification of transposable elements (TE), ortholog inference, and functional annotation particularly in diatoms, which often show high repeat content, dynamic TE landscapes, and lineage-specific gene expansions (Smetacek 1999, Tréguer et al. 2018). Comparing independently produced genome assemblies of the same species provides an opportunity to evaluate structural consistency, identify assembly-specific biases, assess annotation robustness, and quantify functional coherence across workflows. This is particularly valuable for diatoms, where genomic complexity and limited reference genomes continue to challenge comparative and evolutionary analyses (Kooistra et al. 2007).

In this study we compare a resurrected strain (*C. muelleri* BS20) and a lab strain (*C. muelleri* NMCA1316). The resurrected strain has been revived from Baltic Sea sediments from the last century and captures the authentic genetic composition of historical populations, free from the artificial selection, culture drift, and domestication effects that commonly accumulate in long-term laboratory strains. In contrast, a lab strain reflects decades of adaptation to stable culture conditions, often resulting in reduced genetic diversity, altered physiological traits, and the loss of ecologically relevant functions. Sequencing a resurrected strain offers the unique advantage of accessing past genomic diversity and reconstructing evolutionary trajectories over centuries to millennia, enabling direct comparison between ancient and modern populations.

Baltic Sea sediments preserve exceptionally long-lived resting stages of *Chaetoceros muelleri* and other diatoms (for e.g. *Skeletonema marinoi*) (Sanyal et al. 2022, Bolius et al. 2025), providing genetically intact archives that allow researchers to link genomic change to historical environmental shifts such as eutrophication, salinity fluctuations, and climate-driven ecosystem restructuring. Globally, *Chaetoceros* is the most abundant and diverse (58 *Chaetoceros* spp. only in brackish Baltic Sea) marine diatom genus and plays a major role in marine primary production (Malviya et al. 2016, Hällfors 2004). *Chaetoceros* resting spores are found throughout the Baltic Sea stratigraphy from early Littorina Sea (∼7500 cal. yr BP) till date (Andrén et al. 2000) and their distribution correlates with the salinity gradient with higher diversity in the more marine waters (Winter et al. 2010, Hällfors 2004). Hence, *Chaetoceros* will be an ideal system to study diatom evolution over long timescales.

In addition to the two *C. muelleri* strains being resurrected and lab strains, we also present a comprehensive comparative analysis of two independently generated genome assemblies of *C. muelleri*. Our goals are to: (i) disentangle biological versus technical sources of genomic variation by jointly comparing strain-specific differences and assembly-based discrepancies, (ii) identify core genomic features that are consistently represented across both strains and assembly pipelines, (iii) characterize genuine biological divergence between the resurrected and laboratory strains—including structural variation, TE dynamics, gene family differences, and functional annotation profiles—(iv) evaluate assembly- and pipeline-dependent biases affecting repeat resolution, gene prediction, and orthology inference, and (v) establish an integrated genomic framework that incorporates both historical and contemporary diversity to support ecological, evolutionary, and physiological research on this ecologically significant diatom.

## Methods

### Sampling

Sediment sampling was carried out from R/V Electra at Askö in September 2020 using a 1-m gravity corer to recover the uppermost unconsolidated sediment. The detailed sampling protocol has been described in Sanyal et al. (2026).

### Lithology and sample selection

The uppermost section of the core (0–37 cm) consisted of black, gas rich mud with high organic content and no visible laminae. Based on the results from radiometric dating and organic matter measurements (loss on ignition), 12 sediment samples from 0 -32 cm corresponding to 2020 to 1965 CE (Table S1) were selected for the revival of *Chaetoceros* resting spores and subsequent cultivation to obtain high quality DNA.

### Radiometric dating and age modelling

Radiometric dating using 210Pb and 137Cs was performed on sub-samples from the sediment core. Activities of 210Pb, 226Ra, 137Cs and 241Am were measured by direct gamma spectrometry at the Environmental Radioactivity Laboratory, University of Liverpool, using Ortec HPGe GWL-series well-type, low-background intrinsic germanium detectors (Appleby et al., 1986). The results of the radiometric analyses indicate a relatively uniform sedimentation rate since the early 1990s. Sediments below 33 cm are highly compacted and most likely reflect a hiatus in the sediment record, with material below 39 cm being substantially older. Details of the dating methods are provided in Sanyal et al. (2026). The final age–depth model was established following an integrated assessment of all available data, applying the procedures described in Appleby (2001) to determine the most robust chronology.

### Isolation and germination of single resting spores from sediments from the last century

Only the individual germinated resting spores were isolated with care following the protocol of Throndsen (1978). Detailed protocols of the germination and isolation of single resurrected resting spores are provided in Sanyal et al. (2022). Unialgal cultures of *C. muelleri* BS20 derived from single resting spores originating from sediments from the last century were established following the protocol described in Sanyal et al. (2022).

### DNA extraction and sequencing

DNA Isolation from Revived Cultures of Last-Century Diatom Resting Spores. Total DNA from the unialgal *C. muelleri* BS20 culture was extracted using a gentle lysis and phenol–chloroform protocol optimized for diatoms, as described in Sanyal et al. (2022).

### PCR sequencing

PCR amplification and Sanger sequencing were performed to confirm that the taxa is *C. muelleri* using taxonomic markers for centric diatoms used in a previous study by Lee et al. (2013). The primers, PCR amplification and sequencing methodologies are provided in Sanyal et al. (2022).

### Library Preparation and Sequencing

A total of 1,060 ng of extracted DNA was used to prepare a sequencing library with the PacBio SMRTbell prep kit 3.0, following the manufacturer’s instructions (PacBio, Menlo Park, USA). The finished library was sequenced on a PacBio Revio instrument using one SMRT Cell 25M, according to the manufacturer’s recommended protocols. Sequencing was carried out in high-fidelity (HiFi) mode, in which circular consensus sequencing (CCS) was used to generate highly accurate long reads through multiple passes of the polymerase around each SMRTbell template.

### Strains, sequencing, and data overview

Two *C. muelleri* strains (a resurrected strain (*C. muelleri* BS20) and a laboratory strain (*C. muelleri* NMCA1316) were analyzed. Both genomes were generated from PacBio long-read data; BS20 was sequenced with HiFi (CCS) reads (Sanyal et al. 2026), and NMCA1316 with long reads suitable for ABySS hybrid workflows (https://www.ncbi.nlm.nih.gov/datasets/genome/GCA_019693545.1/). Read QC included PacBio’s CCS generation (for BS20) and standard adapter/length filtering (for both datasets).

### Genome assembly, decontamination and completeness

The *C. muelleri* BS20 resurrected strain was assembled with a PacBio HiFi optimized pipeline (hifiasm v0.19.6 (Cheng et al. 2021, 2022) and Flye v2.8.3). The final assembly was generated by careful taxonomic filtering using BLAST (Camacho et al. 2009) and rRNA identification barrnap, https://github.com/tseemann/barrnap) to assign the contigs to the nuclear genome and the non-nuclear contigs to the organelle assembly.

In contrast the long-read PacBio sequence data of the laboratory strain *C. muelleri* NMCA1316 were assembled using ABySS v2.1.5 (de Bruijn graph), and k-mer optimization to maximize contiguity. Assembly statistics (size, N50, GC%, contig count, contig length, Ns/gaps) were computed using the tool ‘assembly-stats’. Completeness was evaluated with BUSCO v5 against eukaryota_odb10 and stramenopiles_odb10.

### Repeat classification, TE class composition and genome masking

A species-specific repeat library was generated with RepeatModeler2 v2.0.3. The library was curated by removing known protein-coding sequences via BLASTP v2.7.1 against the UniProt database, followed by ProtExcluder v1.2. The assembly was masked using RepeatMasker v4.1.2-p1 with the curated library. We annotated repeats with RepeatMasker against the strain specific library and the curated Dfam core HMMs where informative. Genome-wide repeat fraction: proportion of assembly bases overlapping any RepeatMasker annotation (including simple/low-complexity were reported and TE-only (LTR/LINE/SINE/DNA/RC/Unknown) fraction excluding simple repeat and low complexity classes were reported.

### TE class composition and family-level asymmetry

Class-level summaries (e.g., LTR/Copia, LTR/Gypsy, LTR/Ngaro, DNA subclasses) were derived from RepeatMasker annotations. For family-level contrasts, we combined per-family length and copy counts. Shared vs. unique families were defined by exact family identifiers emitted by RepeatModeler2 across the two strain-specific libraries; we note that synonymy among closely related models can inflate uniqueness (addressed in the Discussion/Limitations). We ranked families by Δ masked bp (BS − NMCA1316) to quantify contributors to between-strain differences.

### Gene prediction and structural and functional annotation

Each assembly was annotated using., MAKER v3. annotation pipeline, which integrated evidence-guided training of AUGUSTUS to refine ab initio gene models. To ensure a non-redundant gene set, only one isoform per locus (longest CDS) was retained. Protein models shorter than 50 amino acids or lacking a valid start or stop codon were excluded from downstream analyses. Protein sequences were functionally annotated using eggNOG-mapper v2.1.13 (KEGG orthologs (KO)/COG assignments), and HMMER searches against Pfam-A (gather thresholds). We retained only best KO assignments per gene; Pfam domains were filtered at curated cut-offs to minimize false positives.

### Orthogroup inference and copy-number asymmetry

We inferred orthogroups (OGs) with OrthoFinder v2.5.5 on the predicted proteomes of both strains (DIAMOND mode; default inflation). Across all OGs, shared OGs (genes present in both strains), BS20-unique and NMCA1316-unique OGs were identified. Copy-number comparisons within shared Ogs were estimated from gene counts per strain. We performed Fisher’s exact tests on 2×2 contingency tables per OG (observed counts vs. pooled background), followed by Benjamini–Hochberg (BH) FDR correction. We defined robust expansions as OGs with Δ□≥□2 copies (|BS20 − NMCA1316|□≥□2) and FDR ≤□0.05. To further guard against annotation/fragmentation artifacts, we conducted a permutation test on the Δ□≥□2 subset: gene labels were randomly permuted across strains within each OG (10,000 iterations) while preserving OG sizes; empirical p values were BH-adjusted (FDR_perm), and OGs with FDR_perm ≤□0.05 were retained.

### Functional profiling and enrichment analyses

We summarized KEGG KOs, COG categories, and Pfam domains at the genome level and for the expanded OG gene sets. Fisher’s exact tests with BH correction (FDR ≤□0.05) were done for estimating enrichment. Backgrounds were the annotated genes in each strain for genome-level analysis, and all genes in shared OGs for copy-number analysis.

### TE–gene proximity analysis (strand-aware)

Using the high-contiguity BS20 genome, we assessed the spatial relationship between TEs and genes: Promoter windows were defined strand-aware relative to the annotated Transcription start site (TSS): ≤1□kb promoter (−1,000 to −1 bp upstream of TSS on the sense strand; the complementary downstream interval for genes on the reverse strand) and 1–5□kb promoter (−5,000 to −1,001 bp). Gene-body overlaps were TE intervals intersecting any exon or intron. We computed TE–feature intersections with bedtools intersect (−s for strand where applicable). Genes were assigned to three categories (gene-body; ≤1□kb promoter; 1–5□kb promoter), preferring gene-body > ≤1□kb > 1–5□kb when multiple hits occurred. Impacted gene sets (158 gene body; 60 promoter ≤1□kb; 370 promoter 1–5□kb) were tested for GO/KO/Pfam enrichment against all annotated BS genes as background using Fisher’s exact test with BH correction. For functional context only, we mapped their OGs. to the NMCA1316 annotation and summarized ortholog functions. No TE proximity was computed in the fragmented assembly. All statistical tests visualizations were generated in R (ggplot2) and Python (matplotlib, seaborn).

## RESULTS

### Assembly and annotation of two C. muelleri strains

The two *C. muelleri* genomes (*C. muelleri* BS20, *C. muelleri* NMCA1316) exhibit pronounced differences in assembly contiguity, completeness, and gene annotation quality (Tables 1-3) reflecting both underlying biological divergence and substantial technical variation due to the assembly strategies employed. The assembly metrics demonstrate that the *C. muelleri* BS20 PacBio HiFi assembly provides a substantially more contiguous, better annotation quality and completeness than the ABySS-assembled *C. muelleri* NMCA1316 genome even though both genomes were generated using PacBio long-read sequencing.

**Table 1.** Assembly Metrics for the NCBI Genome vs. In-house Genome.

| Assembly Metric | In-house Genome | NCBI Genome |
| --- | --- | --- |
| Total assembly length (bp) | 43,360,835 | 37,735,205 |
| Number of contigs | 54 | 8,639 |
| Average contig length (bp) | 802,978.43 | 4,368.01 |
| Largest contig (bp) | 3,084,276 | 213,903 |
| N50 (bp) | 1,396,202 | 21,121 |
| N50 contig count | 10 | 452 |
| N60 (bp) | 1,292,182 | 14,242 |
| N60 contig count | 13 | 670 |
| N70 (bp) | 954,505 | 9,224 |
| N70 contig count | 17 | 999 |
| N80 (bp) | 814,299 | 5,415 |
| N80 contig count | 22 | 1,532 |
| N90 (bp) | 599,736 | 2,239 |
| N90 contig count | 28 | 2,576 |
| Minimum contig length (N100) | 10,265 | 200 |
| Total Ns (N count) | 0 | 65,449 |
| Number of gaps | 0 | 904 |

### Assembly contiguity and genome structure

The *C. muelleri* BS20 assembly is highly contiguous, comprising 54 contigs with an N50 of 1.40 Mb and a maximum contig length of 3.08 Mb. Importantly, it contains no gaps or ambiguous bases (Ns), reflecting the resolving power of modern HiFi-based assemblers for repetitive and complex genomic regions. The *C. muelleri* NMCA1316 by comparison, consists of 8,639 contigs, with an N50 of 21.1 kb, a largest contig of 213.9 kb, and 65,449 Ns across 904 gaps. The consequences of the different assemblies are evident in repeat reconstruction. The *C. muelleri* BS20 genome resolves >500 RepeatScout families, including large LTR retrotransposon lineages and numerous high-copy “Unknown” families, whereas the *C. muelleri* NMCA1316 assembly recovers only ∼40–100 families with considerably reduced copy numbers (Table 1). The improved reconstruction of repetitive regions, multi-copy gene families, and regulatory sequences in the *C. muelleri* BS20 genome forms a robust foundation for downstream analyses of functional divergence, including OG structure, metabolic pathways, and domain architecture.

### BUSCO completeness

BUSCO analyses with the lineage-specific stramenopiles_odb10 dataset reveals high completeness for both assemblies (*C. muelleri* BS20: 93.0%, *C. muelleri* NMCA1316: 92.0%). However, resurrected strain *C. muelleri* BS20 shows a higher fraction of duplicated BUSCOs (5 %) compared with the lab strain *C. muelleri* NMCA1316 (1 %), a lower proportion of fragmented BUSCOs (1 %) than the lab strain (2 %), and a lower single-copy BUSCOs (88 %) than the lab strain (91 %) consistent with precise reconstruction of multi-copy gene families and paralogous loci (Table 2). Similarly, both genomes show comparable overall completeness (*C. muelleri* BS20: 72.6%, *C. muelleri* NMCA1316: 71.0**%**) when using the eukaryota_odb10 dataset. These findings further underscore the impact of assembly strategy, and fragmentation patterns on gene recovery. estimation of duplicated gene regions, and single-copy BUSCOs, patterns indicative of assembly artifacts rather than true biological differences.

**Table 2.** Comparative BUSCO summary for the resurrected and laboratory strains of the *Chaetoceros muelleri* genomes using the eukaryota_odb10 and stramenopiles_odb10 datasets.

| BUSCO Metric | Resurrected strain Genome | Lab strain Genome (Protein) |
| --- | --- | --- |
| Dataset: eukaryota_odb10 (n = |  |  |
| Complete BUSCOs (C) | 72.6% (185) | 71.0% (181) |
| □ Complete single-copy (S) | 65.1% (166) | 68.6% (175) |
| □ Complete duplicated (D) | 7.5% (19) | 2.4% (6) |
| Fragmented (F) | 10.6% (27) | 12.9% (33) |
| Missing (M) | 16.8% (43) | 16.1% (41) |
| Total BUSCO groups | 255 | 255 |
| Dataset: stramenopiles_odb10 (n = |  |  |
| Complete BUSCOs (C) | 93.0% (93) | 92.0% (92) |
| □ Complete single-copy (S) | 88.0% (88) | 91.0% (91) |
| □ Complete duplicated (D) | 5.0% (5) | 1.0% (1) |
| Fragmented (F) | 1.0% (1) | 2.0% (2) |
| Missing (M) | 6.0% (6) | 6.0% (6) |
| Total BUSCO groups | 100 | 100 |

### Gene annotation outcomes and structural consequences

The differences in assembly contiguity between the two genomes are directly reflected in their gene annotation statistics (Table 3). The *C. muelleri* BS20 contains 9,723 annotated genes, whereas the *C. muelleri* NMCA1316 genome contains 10,652, a pattern consistent with the overestimation of gene models commonly observed in fragmented assemblies with high contig counts and numerous gaps. Thus, the *C. muelleri* BS20 genome reveals a more structurally coherent and biologically plausible annotation profile. The *C. muelleri* BS20 genome annotations show 23,904 exons and 14,181 introns, compared with 25,179 exons and 14,527 introns in the lab strain. The increased exon and intron count in the lab strain likely arise from contig breakpoints splitting multi-exon genes, creating artificially fragmented transcripts and increasing the number of reconstructed gene models. The higher number of single-exon mRNAs in the lab strain (3,484 vs. 2,715 in *C. muelleri* BS20) further supports the overestimation consistent with partial ORFs generated at scaffold ends. Furthermore, despite low absolute numbers due to stringent UTR prediction, the *C. muelleri* BS20 genome contains 1–2 transcripts with annotated 5′ or 3′ UTRs, compared to 2–3 in the lab strain. the increase in UTR counts in the lab strain likely reflects false positives associated with broken transcript models due to fragmentation rather than improved completeness.

**Table 3.** Genomic features of the resurrected genome vs. the lab strain.

| Genomic | Resurrected strain | Lab strain |
| --- | --- | --- |
| Number of gene | 9723 | 10652 |
| Number of mrna | 9723 | 10652 |
| Number of mrnas with utr both sides | 1 | 2 |
| Number of mrnas with at least one utr | 1 | 3 |
| Number of cds | 9723 | 10652 |
| Number of exon | 23904 | 25179 |
| Number of five_prime_utr | 1 | 2 |
| Number of three_prime_utr | 1 | 3 |
| Number of exon in cds | 23904 | 25179 |
| Number of exon in five_prime_utr | 1 | 2 |
| Number of exon in three_prime_utr | 1 | 3 |
| Number of intron in cds | 14181 | 14527 |
| Number of intron in exon | 14181 | 14527 |
| Number gene overlapping | 5 | 0 |
| Number of single exon gene | 2715 | 3484 |
| Number of single exon mrna | 2715 | 3484 |

### Orthogroup Divergence Between Strains

#### Orthogroups

A total of 27,815 OGs was identified, 7,196 OGs were shared between strains, with 416 unique to *C. muelleri* BS20 and 748 unique to the lab strain. Among shared OGs, 538 show higher copy number in the resurrected *C. muelleri* BS20 strain and 439 enriched in the lab strain. A Fisher exact test followed by Benjamini–Hochberg correction, shared confirmed that these copy number expansions within these shared OGs (538 and 439 OGs) are statistically significant (FDR ≤ 0.05). A total of 6,219 of the shared 7,196 OGs revealed highly conserved equal copy number consistent with broad stability of gene family size across lineages irrespective of differences in genome contiguity. In total, 9,437 *C. muelleri* BS20 and 10,218 *C. muelleri* NMCA1316 genes are assigned to OGs (8,872 and 8,909 in shared OGs), indicating more OG-assigned models in the lab strain annotation but with modest copy-number asymmetries that align with assembly and annotation differences. To distinguish robust biological expansions from noise introduced by annotation or fragmentation, we applied a Δ□≥□2 copy threshold, identifying 53 OGs with higher copy number in the resurrected strain and 80 in the lab strain. A permutation-based test applied to the Δ□≥□2 subset validated these robust expansions, with all 53 and 80 OGs remaining significant at FDR_perm ≤ 0.05 (Fig. 1).

**Fig. 1.**
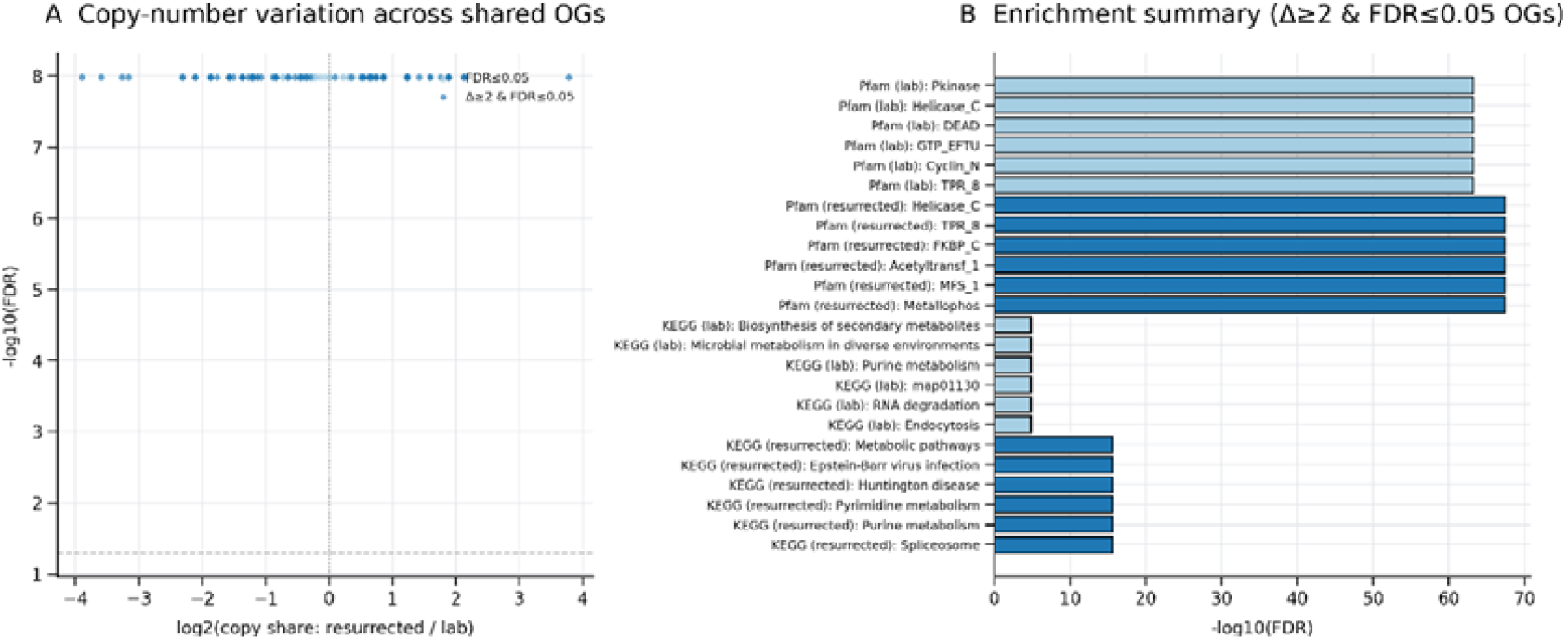
(A) Volcano plot showing log□(copy-share: resurrected/lab) vs log□□(FDR) from per-OG Fisher tests (BH correction). Points in dark blue are Δ≥2 & FDR ≤ 0.05 (robust expansions); light blue are FDR ≤ 0.05 but Δ<2; grey are not significant. (B) Enrichment summary of top KEGG pathways and Pfam domains enriched in Δ≥2 & FDR ≤ 0.05 OGs for each strain; bars show log□□(FDR); blue = resurrected; light blue = laboratory strain.

### Functional identity of expanded orthogroups

Functional comparisons between the two *C. muelleri* strains reveal a large, shared core functional repertoire, alongside smaller strain-specific gene sets. In total, 2,517 KOs were shared between the strains, with 288 unique to the resurrected BS20 strain and 392 unique to the laboratory strain. Similarly, 2,934 Pfam domains were shared, with 166 domains unique to BS20 and 261 unique to the laboratory strain. In contrast, COG functional categories showed complete overlap, with all 24 COG classes represented in both strains, indicating broad conservation of major cellular systems despite differences in individual gene content.

We examined copy-number variation focused on OGs showing Δ≥2 gene copy differences between strains. At this level, KEGG and Pfam annotations revealed contrasting functional biases between the two assemblies, whereas COG category representation remained largely similar. These OG-level differences mirror patterns observed in the strain-specific KEGG and Pfam repertoires and suggest a combination of biological variation and assembly-driven annotation effects, particularly in regions rich in repetitive or low-complexity sequence.

In the resurrected *C. muelleri* BS20 strain, Δ≥2 expanded OGs comprised 417 genes, compared against a background of 6,518 genes from shared OGs. KEGG enrichment analysis identified 14 significantly enriched KOs (FDR ≤ 0.05), including K13993, K06269, K12863, and related orthologs associated with DNA and RNA processing, replication-linked functions, and lineage-specific enzymatic roles, several with extremely strong statistical support (p□<□1□×□10^−1^□ and high odds ratios).

Pfam enrichment revealed a broader signature, with 48 significantly enriched domains, many corresponding to retrotransposon-associated families (e.g., gag– pre-integrase regions and gag-like retrotransposon domains), along with nucleic-acid-binding domains and stress-associated elements such as HSP20. Together, these patterns indicate that gene family expansions in BS20 disproportionately involve TE-rich genomic regions and functions associated with genomic regulation and stress responsiveness, consistent with the improved recovery of such regions in the long-read HiFi assembly (Fig. 1).

In contrast, Δ≥2 expanded OGs in the laboratory strain comprised 551 genes, relative to 6,511 genes from shared OGs. KEGG analysis identified 22 significantly enriched KOs, including K07374, K07820, and K00820, with prominent signals related to carbohydrate metabolism, glycosyltransferase activity, and lineage-specific catalytic functions (many with p□<□1□×□10^−^□). Pfam enrichment shows 104 significantly enriched domains, including plant associated transmembrane domains, galactosyltransferase domains, LCCL and NHL repeat domains associated with protein–protein interaction scaffolding, multiple HTH/DDE DNA transposase associated domains as well as several retroelement and virus-derived domains. While some enriched categories are plausibly linked to cellular processes such as polysaccharide metabolism or membrane trafficking, the higher degree of fragmentation in the laboratory strain assembly suggests that inflated gene counts arising from split or partial ORFs may contribute substantially to these patterns. Thus, functional enrichments inferred from the laboratory strain should be interpreted cautiously, primarily reflecting the combined effects of biological variation and assembly quality (Fig. 1).

Overall, KEGG, COG, and Pfam analyses demonstrate that the two *C. muelleri* strains share a broadly conserved functional architecture, while differing in gene family composition at finer scales. Importantly, robust functional enrichment signals can be confidently interpreted only for the high-contiguity BS20 assembly, whereas apparent expansions in the laboratory strain underscore the strong influence of assembly fragmentation on downstream functional inference.

### Repetitive DNA landscape in two C. muelleri strains

#### Genome–wide repeat fractions

Using non-overlapping (union) intervals, the resurrected BS20 strain contains 16,710,566□bp of repeats in a 44,084,192□bp assembly, i.e. 37.91□% of the genome is repetitive. The laboratory strain has 8,046,829□bp of repeats in a 37.7□Mb assembly, i.e. 21.34□% repetitive. Thus, the resurrected genome harbors ∼1.8-fold repetitive DNA than the laboratory strain.

#### Class composition and family-level asymmetry

Examining the TE composition in the two strains revealed that in both the strains, the majority of masked TE bases were assigned to Unknown categories—8.30□Mb (51.9%) in the resurrected genome versus 3.55□Mb (47.6%) in the laboratory strain which is consistent with many novel, currently unclassified repeat families identified by RepeatModeler. Beyond “Unknown,” the resurrected strain was enriched for LTR/Copia (3.83□Mb; 23.9%) and unresolved LTR (1.13□Mb; 7.08%), as well as several DNA transposon subclasses (PIF-Harbinger: 0.374□Mb; 2.34%; MULE-MuDR: 0.231□Mb; 1.44%; PiggyBac: 0.133□Mb; 0.83%). In contrast, the laboratory strain showed relatively larger fractions of LTR/Gypsy (1.48□Mb; 19.9% vs 1.35□Mb; 8.46% resurrected strain) and LTR/Ngaro (0.298□Mb; 4.00% vs 0.103□Mb; 0.64% resurrected strain). Overall, TE content (excluding microsatellites/low-complexity) in the resurrected strain was 15.99□Mb versus 7.46□Mb in the laboratory strain (Δ□=□8.52□Mb). The largest contributors included one LTR/Gypsy family (1.00□Mb), a generic LTR family (0.997□Mb), four Unknown families (0.577, 0.447, 0.427, 0.395□Mb), and three LTR/Copia families (0.550, 0.445, 0.273□Mb) (Table 4). At the family level, the overrepresentation in the resurrected strain was concentrated in a small number of novel families: the top 7 families (Table 5) in the resurrected strain only explained ∼52.2% (4.45□Mb) of the between-strain difference, and the top 18 families explained ∼80.0% (6.82□Mb).

**Table 4:** Comparative TE content between the resurrected and the laboratory strains.

| TE Class / Family | In-house | In-house | Reference | Reference | Enriched In |
| --- | --- | --- | --- | --- | --- |
| Unknown | 51.9% | 8.30 Mb | 47.6% | 3.55 Mb | Resurrected |
| LTR/Copia | 23.9% | 3.83 Mb | 19.2% | 1.43 Mb | Resurrected |
| LTR (unresolved) | 7.08% | 1.13 Mb | 0.062% | 4.6 kb | Resurrected |
| LTR/Gypsy | 8.46% | 1.35 Mb | 19.9% | 1.48 Mb | Laboratory |
| LTR/Ngaro | 0.64% | 0.103 Mb | 4.00% | 0.298 Mb | Laboratory |
| DNA / | 2.34% | — | 0.62% | — | Resurrected |
| DNA / MULE-MuDR | 1.44% | — | 0.68% | — | Resurrected |
| DNA / PiggyBac | 0.83% | — | 0.62% | — | Resurrected |
| Total TE (TE-only) | — | 15.99 Mb | — | 7.46 Mb | Resurrected ($\Delta=8.52$ |

**Table 5.**
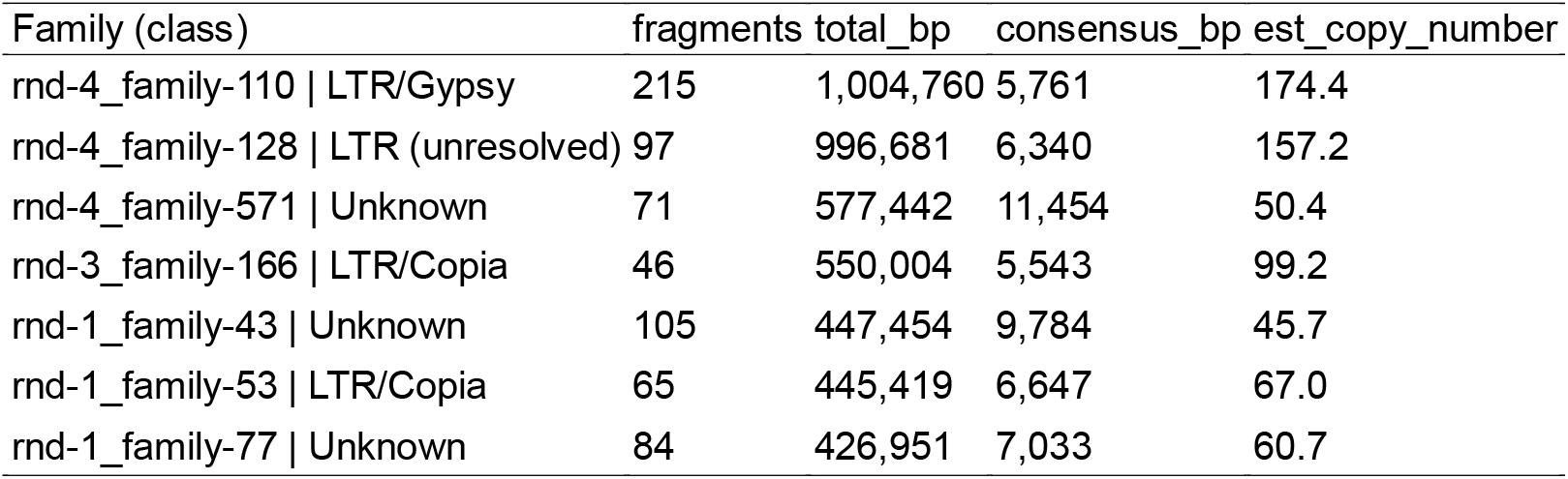
The overrepresentation of the TE families exclusively in the resurrected strain.

| Family (class) | fragments | total_bp | consensus_bp | est_copy_number |
| --- | --- | --- | --- | --- |
| rnd-4_family-110 \| LTR/Gypsy | 215 | 1,004,760 | 5,761 | 174.4 |
| rnd-4_family-128 \| LTR (unresolved) | 97 | 996,681 | 6,340 | 157.2 |
| rnd-4_family-571 \| Unknown | 71 | 577,442 | 11,454 | 50.4 |
| rnd-3_family-166 \| LTR/Copia | 46 | 550,004 | 5,543 | 99.2 |
| rnd-1_family-43 \| Unknown | 105 | 447,454 | 9,784 | 45.7 |
| rnd-1_family-53 \| LTR/Copia | 65 | 445,419 | 6,647 | 67.0 |
| rnd-1_family-77 \| Unknown | 84 | 426,951 | 7,033 | 60.7 |

Using strand-aware annotation in the high-quality resurrected genome, our TE-gene proximity analysis identified 158 genes with TE insertions within gene bodies, 60 genes with TE insertions within ≤1 kb upstream promoters, and 370 genes with TE insertions within 1–5 kb promoters. Enrichment tests applied to the TE-impacted gene sets in the resurrected genome did not yield significant GO, PFAM-A, or KEGG KO terms after FDR correction, consistent with small gene lists and sparse annotations. To examine the functional identity of these genes, the OGs of the TE-impacted genes in the resurrected genome were also evaluated in the lab strain. This homolog-based analysis revealed strong enrichment for transmembrane transport and membrane-associated functions (e.g., transporter activity GO:0022857, transmembrane transport GO:0055085, ion transport), but importantly these enrichments reflect conserved functions of the gene families impacted in the resurrected genome—not TE activity in the fragmented lab strain assembly, where TE–gene spatial relationships were not assessed.

Furthermore, 2,411 TE families were shared across the two strains, 2,667 and 3,129 families exclusive to the resurrected and the laboratory strain were identified (Table□4). These “unique” family counts should not be interpreted as truly lineage-specific TE innovations, because TE family identification is sensitive to sequence fragmentation, consensus length, and annotation redundancy. However, even after accounting for these annotation artefacts, the large asymmetry in non-shared families strongly suggests genuine TE turnover between the two genomes, consistent with independent TE expansions, differential retention, and family-specific decay processes.

## Discussion

### Assembly strategy strongly determines genome contiguity and gene-space recovery

Our comparative analysis of two *C. muelleri* strains demonstrates that assembly strategy is a primary determinant of genome quality, even when both genomes are derived from PacBio long-read sequencing. The resurrected *C. muelleri* BS20 strain, assembled using HiFi-optimized pipelines (hifiasm and Flye), achieved high contiguity, comprising 54 contigs with an N50 of 1.40□Mb, a maximum contig length of 3.08□Mb, and no gaps (0 Ns), yielding a highly contiguous, near-gapless representation of the nuclear genome. In contrast, the laboratory strain NCMA1316 exhibited substantial fragmentation, with 8,639 contigs, an N50 of 21.1□kb, a largest contig of 213.9□kb, and 65,449 Ns across 904 gaps (Table□1).

These differences in contiguity had pronounced effects on downstream repeat and gene annotation. The BS20 assembly enabled reconstruction of >500 RepeatScout families, including large LTR retrotransposon lineages and numerous high-copy “Unknown” families, accounting for 16.71□Mb of repetitive DNA (37.91□%), together with accurate recovery of 7.5□% duplicated BUSCOs. In contrast, repeat recovery in the laboratory strain was markedly reduced (8.05□Mb; 21.34□%), with only ∼40–100 repeat families, substantially lower copy numbers, and fewer duplicated BUSCOs (2.4□%). Gene counts were correspondingly inflated in the laboratory strain (10,652 vs. 9,723 in BS20), consistent with open reading frame (ORF) splitting driven by assembly fragmentation, particularly in repeat-rich and gene-dense regions.

These patterns indicate that the observed disparities primarily reflect differences in assembly resolution rather than true biological absence of repeats or gene families in the laboratory strain. The superior contiguity of the BS20 assembly provides more accurate representation of gene families embedded within TE-rich genomic regions, whereas fragmentation in the NCMA1316 assembly leads to collapsed repeats, truncated consensus models, and inflated gene counts (Table□2). Thus, these results highlight a central challenge in microbial eukaryote genomics: biological signal can be obscured by a combination of assembly strategy, raw read data, and upstream DNA quality, even when long-read sequencing is employed. Thus, our findings underscore the importance of HiFi-optimized pipelines and high-quality input DNA for resolving repetitive architecture and for robust inference of gene family evolution, regulatory complexity, and genomic innovation in diatoms and other TE-rich microalgae.

### Biological differences between strains are superimposed on assembly-driven variation

Although assembly quality explains many of the discrepancies between the two genomes, genuine biological differences between strains were also evident. The resurrected strain BS20 originates from Baltic Sea sediments deposited in the last century, whereas NMCA1316 is a contemporary laboratory strain propagated under stable conditions for years or decades, exposing the two strains to markedly different environmental and selective regimes.

The two strains share 7,196 OGs, with 6,219 showing identical copy numbers, and exhibit near-identical COG profiles, with all 24 COG categories present in both genomes, indicating a strongly conserved genomic backbone. Functional annotation further revealed 2,517 shared KOs and 2,934 shared Pfam domains, and BUSCO completeness was similarly high in both assemblies (93.0□% in BS20 and 92.0□% in NMCA1316, using *stramenopiles_odb10*). Thus, these shared features demonstrate that the core metabolic, regulatory, and cellular functions of *C. muelleri* are conserved across strains and robust to differences in assembly methodology.

A total of 166 Pfam domains were unique to BS20 and 261 to the laboratory strain, these differences should be interpreted with caution. In particular, the higher number of strain-specific Pfam domains observed in NMCA1316 is likely influenced by assembly fragmentation and annotation inflation, rather than reflecting distinct evolutionary histories. Similarly, Pfam domain uniqueness in the laboratory strain primarily highlights the impact of assembly quality on domain detection and gene model completeness, whereas functional enrichments and gene family expansions inferred from the highly contiguous BS20 assembly provide a more reliable basis for biological interpretation.

After applying stringent Δ≥2 copy thresholds and permutation-based significance testing, 53 OG families were robustly expanded in the resurrected *C. muelleri* BS20 strain, compared with 80 expanded OGs in the laboratory strain NMCA1316. The functional signatures of these expansions differed considerably between strains.

In BS20, expanded OGs were enriched for retroelement associated domains, nucleic acid binding and processing proteins, and stress responsive gene families, including small heat shock proteins (sHSPs) such as HSP20. Small HSPs are widely implicated in tolerance to thermal, oxidative, osmotic, and metabolic stress (Huang et□al.□2025). In diatoms, heat-shock pathways—including sHSP-associated genes—contribute to thermal tolerance and antioxidant defense, with heat-shock transcription factor cascades activating stress-response genes under elevated temperatures (Huang et□al.□2025). Because sHSPs function independently of ATP, they remain active under conditions of limited cellular energy availability (Sato et□al.,□2024), making them particularly relevant during low-energy physiological states such as prolonged darkness, hypoxia, or sediment burial. Although direct evidence in diatom resting spores is still emerging, sHSPs including HSP20 are upregulated during life-cycle transitions and resting-stage formation in other microalgae, such as the dinoflagellate *Scrippsiella trochoidea*, indicating a conserved role in dormancy-associated stress tolerance (Deng et□al.□2020).

In contrast, OGs expanded in the laboratory strain were enriched for glycosyltransferases, carbohydrate-modifying enzymes, membrane-trafficking modules, and several DNA transposase associated domains. While some of these enrichments may reflect metabolic specialization under long-term culture conditions, they are also likely influenced by assembly fragmentation, which can inflate gene counts through ORF splitting and partial gene models.

Consistent with this interpretation, the BS20 genome retained a broader and more coherent repertoire of TE-associated gene families, including gag-like and retrotransposon-derived domains, together with stress-responsive genes such as HSP20 and other nucleic acid binding factors. Many of these genes were located within or proximal to TE insertions (158 intragenic and 430 promoter-proximal cases), suggesting that historical environmental variability in the Baltic Sea, combined with an intact TE landscape, has shaped regulatory and stress-associated gene family architectures in the resurrected lineage. By contrast, apparent expansions in the laboratory strain likely reflect a mixture of culture associated divergence and technical artefacts, underscoring the need for caution when interpreting functional novelty from fragmented assemblies.

Our findings reinforce the importance of the assembly quality in comparative genomics, as biological inference can be confounded when fragmentation overestimates gene family expansion. Robust functional signals are most confidently interpreted from the high-contiguity BS20 assembly, highlighting the value of HiFi-optimized approaches for resolving TE-rich and stress-responsive genomic regions in diatoms.

### Transposable element dynamics differentiate historical and modern genomes

One of the most striking differences between the two strains lies in their TE landscapes. The resurrected genome contains 37.9% repetitive DNA, approximately1.8-fold higher than the laboratory strain (21.3%). Much of this difference arises from LTR/Copia elements, unresolved LTRs, and several subclasses of DNA transposons which are robustly represented in long, intact contigs in BS20. In contrast, the laboratory strain shows relatively higher representation of LTR/Gypsy and Ngaro elements, although at reduced absolute abundance.

The expanded TE content in BS20 likely reflects several processes. These include long-term accumulation of TEs in historical populations, potentially maintained through resting stages exposed to fluctuating environmental pressures; superior assembly resolution, enabling accurate reconstruction of high-copy and nested elements; and possible reduction of TE load in the laboratory strain through culture driven mutation, bottlenecks, purging of mobile elements, or sustained propagation under stable conditions. Notably, TE family asymmetry in BS20 was concentrated in a limited number of high-copy families, consistent with episodic TE bursts in the historical lineage.

Analysis of TE–gene proximity further revealed that hundreds of genes in the resurrected strain contain TE insertions within gene bodies or proximal regulatory regions. Although pathway level enrichment did not remain significant after FDR correction, homology-based evaluation in both strains identified recurrent associations with membrane transport and ion translocation related genes, suggesting that TE insertions may preferentially affect loci involved in environmental responsiveness.

Our findings indicate that TE dynamics are a major contributor to genomic divergence between resurrected and laboratory strains, providing insight into how diatom genomes evolve under natural versus long-term culture conditions. The findings highlight potential consequences for regulatory flexibility, stress responsiveness, and long-term adaptive capacity, while also emphasizing the importance of assembly quality for resolving TE-rich genomic regions.

### Resurrected genomes offer unique access to historical diversity and evolutionary trajectories

Our study highlights the exceptional value of resurrected diatom strains for evolutionary genomics. Resting stages recovered from Baltic Sea sediments provide access to historical genomic states that are largely unaffected by long-term laboratory propagation, including domestication, culture drift, and mutational bottlenecks that can reshape modern lab strains. By sequencing BS20, we captured genomic diversity that reflects past environmental regimes, including periods of salinity fluctuation, eutrophication, industrialization, and climate-driven restructuring in the Baltic Sea ecosystem. Comparison with a contemporary laboratory strain reveals how diatom genomes diverge across ecological time scales and under contrasting selective environments. Such comparisons enable reconstruction of TE expansion, gene family turnover, and shifts in metabolic and regulatory repertoires through time, while providing a framework for linking genomic change to environmental histories archived in sediment cores. Thus, these approaches establish a powerful foundation for integrating genomics with paleobiology and long-term ecosystem monitoring.

### Toward a standardized genomic framework for Chaetoceros and other diatoms

Examining two independently assembled genomes from distinct *C. muelleri* strains provides a clear roadmap for future diatom genomics. Our results highlight several key considerations. First, HiFi based long-read assembly pipelines are essential for accurate reconstruction of TE-rich diatom genomes. Second, annotation inflation in fragmented assemblies can substantially mislead functional and comparative analyses. Third, strain history matters: resurrected and long-term laboratory strains represent distinct evolutionary trajectories shaped by contrasting selective regimes. Fourth, OG and analyses of repeats require careful interpretation, especially when assembly contiguity differs between genomes. Finally, integrating historical (resurrected) and modern genomes offers unique opportunities to reconstruct diatom evolutionary trajectories and to link genomic change with ecological and environmental adaptation.

In conclusion, our comparative approach establishes a foundation for a robust, strain-aware genomic framework for *C. muelleri*. Such a framework is essential for interpreting evolutionary change, ecological responses, and functional innovation in diatoms; one of Earth’s most important and diverse groups of microalgae.

## Data availability

All sequencing data and genome assemblies generated in this study are publicly available. The *Chaetoceros muelleri* BS20 genome assembly, raw long-read and short-read sequencing datasets, and RepeatMasker annotations will be deposited in the NCBI Sequence Read Archive (SRA) and NCBI Genome. The previously published *C. muelleri* reference genome used for comparative analyses is available through NCBI under accession GCA_ _019693545.1. Scripts used for genome assembly are archived on GitHub at https://github.com/tseemann/barrnap). Any additional data supporting the findings of this study are provided within the article or are available from the corresponding author upon reasonable request.

## Author contributions

Anushree Sanyal designed the experiment, generated and analyzed the data and wrote the manuscript.

## Acknowledgements

We thank the captain and crew of R/V Electra af Askö for their excellent assistance with coring operations despite pandemic-related restrictions and challenging weather conditions. We gratefully acknowledge Prof. P.□G. Appleby and G.-T. Piliposian (University of Liverpool) for the rapid processing of the radiometric dating analyses. We thank Anna Vilaplana Burgos for her assistance with the DNA extraction. Funding was provided through the REVIVE project, supported by the Foundation for Baltic and East European Studies (grant no.□42-19). This work was supported by the National Bioinformatics Infrastructure Sweden (NBIS) at SciLifeLab. The computations were performed on resources provided by the Swedish National Infrastructure for Computing (SNIC), partially funded by the Swedish Research Council through access to the Dardel supercomputer at the PDC Center for High Performance Computing, KTH Royal Institute of Technology under Project SNIC 2022-22-9. PacBio sequencing was performed by the National Genomics Infrastructure (NGI) Uppsala, SciLifeLab, Sweden.

